# X-ray fluorescence microscopy exposure estimates using a single excitation energy

**DOI:** 10.64898/2026.01.15.699668

**Authors:** Benjamin Roter, Andrew M. Crawford, Thomas V. O’Halloran, Chris Jacobsen

**Affiliations:** Applied Physics Program, Northwestern University, Evanston, IL 60208, USA; Department of Microbiology, Genetics, & Immunology, Michigan State University, East Lansing, MI 48824, USA; Department of Chemistry, Michigan State University, East Lansing, MI 48824, USA; Elemental Health Institute, Michigan State University, East Lansing, MI 48824, USA; Department of Physics and Astronomy, Northwestern University, Evanston, IL 60208, USA; Chemistry of Life Processes Institute, Northwestern University, Evanston, IL 60208, USA

## Abstract

Scanning fluorescence x-ray microscopy is widely used for quantitative mapping of elemental concentrations, including in studies of essential, but low-concentration metals in cells, tissues, and organs. Practical studies often use a single incident photon energy to excite fluorescence from many elements. We present calculations of the number of incident photons per pixel required to detect a specified areal concentration of an element in the case of non-resonant excitation, along with the calculated radiation dose consequently imparted in a simple model tissue. We also show how certain approximations can lead to less accurate estimates. These results can be used to guide experimental planning for studies of the role of low-concentration elements in biological tissues.

## 1. Introduction

X-ray fluorescence (XRF) allows for the identification of specific chemical elements, as was understood more than a century ago [1–3]. Scanning a small x-ray beam allows for the imaging of elemental content [4–6] in an approach which we refer to here as scanning fluorescence x-ray microscopy (SFXM). This approach shows wide utility including in research of the roles of essential, but low-concentration metals in biological functions in 2D [7–9], and in 3D via x-ray fluorescence tomography [10].

In transmission x-ray microscopy using absorption and phase contrast, there is rich literature on calculations of x-ray fluence requirements for achieving a specified spatial resolution [11–15], and similar calculations exist for imaging based on coherent scattering [16, 17]. However, while there have been careful studies of the achieved elemental detection limits in specific synchrotron-based SFXM measurements [18, 19], there have been fewer equivalent calculations aimed at predicting the illumination required to achieve a specified level of detection. Several early studies used approximate values for the relevant interaction coefficients to compare x-ray-induced x-ray fluorescence against other elemental detection methods such as electron- or proton-induced x-ray emission, and electron energy-loss spectroscopy [20, 21]. This methodology was also used to predict x-ray illumination requirements for SFXM [22]. However, these studies assumed illumination at a photon energy just above the relevant absorption edge of each element.

Today, SFXM is often carried out using a single incident photon energy (often 10–12 keV in the case of many biological studies) to excite the emission of x-ray fluorescence from multiple elements simultaneously, where these elements have absorption edges and emission lines at photon energies well below the illumination photon energy. Photon counting at energies characteristic of the elements of interest, coupled with spectrum analysis, background correction, and mass calibration standardization, leads to quantitative elemental maps of the sample. For biologists interrogating tissue and cell-based samples, the pixel-by pixel quantitative resolution of heterogeneity in these elemental maps provides powerful insights into fundamental biological processes, as well as etiology of disease states [23]. To model this common practice and better understand limitations in the quantitative results, one must account for non-resonant excitation.

In most SFXM experiments, energy-dispersive spectrometry (EDS) detectors are used in conjunction with analysis programs [24–27] that account for the energy resolution of such detectors, and backgrounds including x-ray scattering and incomplete charge collection from the detector [28]. Fluorescence spectrum analysis is simplified if one instead uses wavelength dispersive spectrometry (WDS) detectors [29], but WDS is limited in solid angle coverage and wavelength range so that it is usually not employed unless chemical state information is required.

With these developments, we revisit the question of illumination requirements for SFXM imaging of elemental concentrations. We account for both resonant and non-resonant illumination photon energies, and we make use of easy computer access to accurate tabulations of the relevant x-ray interaction coefficients as is described in Sec. 2. This allows us to predict the incident fluence 𝔉 (given here in photons/cm^2^ – see Sec. 3.2) at a single incident photon energy *E*_inc_ required to detect specific elements at a specific areal mass concentration *ρ*^′^. Such calculations are especially interesting when considering a specified minimal value 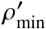, which sets the limit of detection (LOD) for an element. We can then use this fluence 𝔉 to calculate the radiation skin dose *D*_skin_ necessarily imparted to the incident-beam-facing surface of a specified “matrix” material (for example, for detection of an element located in a biological cell with some average composition), since dose can set limits on the imaging of radiation-sensitive materials. We show *xraylib*-based [30] calculations for a wide range of trace elements at different incident energies using an x-ray fluorescence forward model that considers the excitation dependence of mass photoionization cross sections, Coster-Kronig transitions, cascade effects, the presence of an EDS detector entrance window, and the presence of any gas in the sample environment.

## 2. The x-ray fluorescence forward model

In a typical SFXM experiment, the specimen is meant to be illuminated with 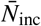 incident photons per pixel at energy *E*_inc_ (per-pixel statistical fluctuations will be distributed around 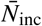). Some fraction of the incident photons are absorbed by a target element of atomic number *Z*, leading ultimately to a mean number of detected photons of 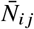corresponding to x-ray fluorescence line *i j*, where *i* and *j* are initial and the final electron vacancy states, respectively. (For notational simplicity, we do not explicitly indicate the energy dependence of each variable, but this is shown in Table 1; see also Sec. S1 in the Supplement for information on relating transitions *i j* to conventional x-ray nomenclature). For a specimen sufficiently thin that there is neither scattering of the incident signal nor self-absorption of the fluorescence signal, the mean number 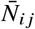 of detected fluorescence photons can be expressed as [5, 20, 30, 31]

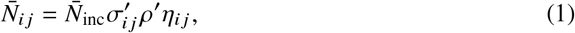

where 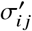 is the mass x-ray fluorescence production cross section (*e*.*g*., cm^2^/g) at energy *E*_inc_, *ρ*^′^ is the local areal mass density (*e*.*g*., g/cm^2^), and η_*i j*_ is the net detection efficiency for line *i j* (see Eq. 5).

**Table 1.**
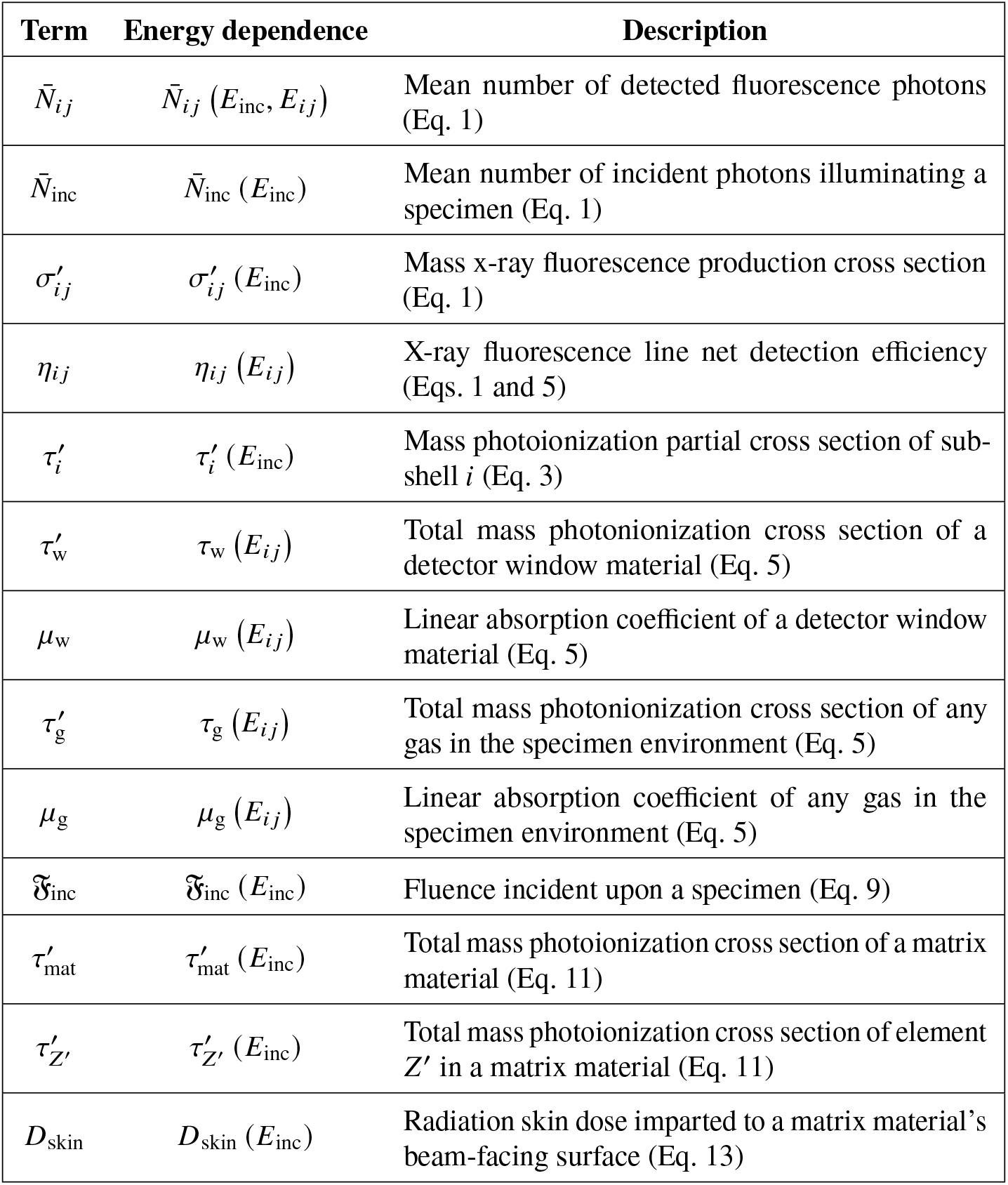
Energy-dependent parameters used in our calculations, with a short description and an indication of where they first appear.

In this work, we use mass cross sections 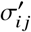 instead of atomic cross sections σ_*i j*_, which are typically given in barns (1 barn = 10^−24^ cm^2^). Mass cross sections are more common to find in fundamental parameter databases [32], but they are related to atomic cross sections via

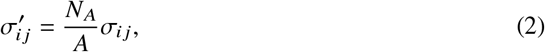

where σ_*i j*_ is the atomic XRF production cross section, *N*_*A*_ is Avogadro”s number, and *A* is the molar mass of the absorbing element.

Mass XRF production cross sections are related to the probability of fluorescence line *i j* being emitted due to subshell *i* being excited by an incident x-ray photon; this can be calculated via [30]

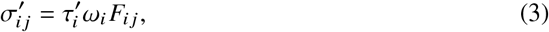

where 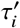 is the mass photoionization partial cross section (PCS) of subshell *i, ω*_*i*_ is the subshell fluorescence yield, and *F*_*i j*_ is the fractional yield or branching ratio which is the fraction of *ω*_*i*_ emitted fluorescence photons corresponding to the *i j* line. For 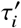, care is generally taken when determining what values to use. The simplest assumption is that as one crosses the threshold energy for removing an electron from a specific subshell, the fractional increase in absorption tells one the fractional increase in fluorescence events resulting from that subshell. This gives rise to the jump ratio approximation *r*_*i*_ for subshell *i* [31, 33, 34]. This approximation generally holds well for *K* shell excitations, with the fraction of absorption events that go towards creating *K* shell vacancies given by

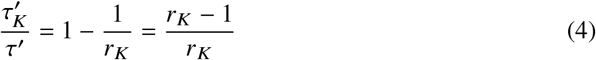

where 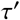 is the total mass photoionization cross section. For the *L* shell and beyond, the excitation-dependent nature of photoionization becomes more important at higher *E*_inc_ relative to the respective subshell absorption edge [35–38]. Therefore, the jump ratio approximation of Eq. 4 loses accuracy when applied to *L* shell fluorescence, as illustrated in Sec. 4. Two additional phenomena [30, 39] play important roles for those shells:

1. Coster-Kronig (CK) transitions: Special cases of Auger electron emission where electron vacancies are filled by electrons in higher subshells within the same shell, causing electrons to be emitted from either higher shells or from the same shell. In the latter case, the CK transition becomes a super Coster-Kronig (SCK) transition.
2. Cascade effects: Vacancies created due to general Auger emission, electrons emitted due to (S)CK transitions, and/or XRF events involving lower shells.

Together, these phenomena can affect the values of 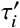, especially as *E*_inc_ goes well beyond edge energies 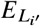 [where *i*^′^ = 1, 2, 3; (similar considerations apply to *M* edges and beyond)]; this is illustrated in calculations shown in Sec. 3.1.

Detection efficiencies η for x-ray fluorescence typically consider the solid angle fraction Ω / (4π) that a fluorescence detector subtends. Most hard x-ray fluorescence experiments use energy-dispersive detectors (EDS), and in most cases the cooled detection elements are protected from contamination by being placed behind thin windows. These windows of thickness *t*_w_ are often fabricated of beryllium so as to minimize the absorption of fluorescence at energies of a few keV or above. In addition to detector entrance windows, we also account for signal absorption in a gas path (air, helium, etc.) over a distance *t*_g_ between the sample and detector window. Therefore, we modify the efficiency η to include window and gas attenuation factors, giving η_*i j*_ for a particular fluorescence line *i j* of

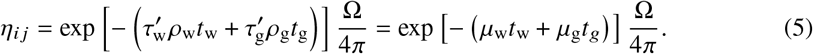

In the above equation, 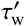 and 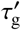 are the total mass photoionization cross sections of the window material and gas, respectively, *E*_*i j*_ is the fluorescence energy of line *i j* of an emitting element, and *ρ*_w_ and *ρ*_g_ are the densities of the window material and gas, respectively. (The second form of Eq. 5 use the energy-dependent window and gas material linear absorption coefficients 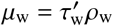 and 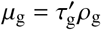, respectively.) We show in Sec. S2 of Supplement 1 how a beryllium window and air can change the minimum number of incident photons 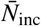 per pixel required to detect a given number of fluorescence photons per pixel at a specified areal mass concentration 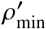. Incomplete charge collection [28] of electron-hole separation events in the detector can also effectively reduce the number of detected photons; however, to simplify calculations, we ignore this factor.

## 3. Calculations of minimum photon exposure and radiation dose

We now use Eq. 1 to solve for the required number of incident photons 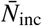 per area. We do so based on a requirement to detect 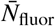 fluorescence photons per pixel in an image.

### 3.1. Minimum number of incident photons per pixel

To obtain the theoretical minimum number of incident photons per pixel 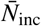 for detecting 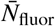 fluorescent photons per pixel, all fluorescence line contributions calculated using Eq. 1 can be summed up via

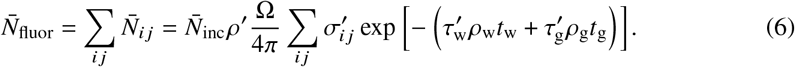

From basic considerations of false-positive and false-negative error rates in detection [40], it is often sufficient to detect 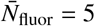 photons in order to detect the presence of an element at low concentration when the background is sufficiently low. With a specified requirement for 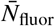, one can rearrange Eq. 6 to solve for the required number of incident photons per pixel 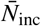. We do so at a specified minimum value of detectable mass density 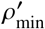 corresponding to a desired limit of detection (LOD); this gives

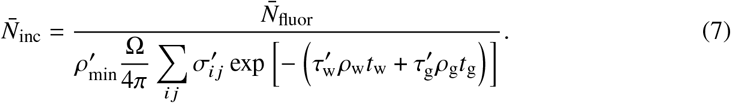

This result allows us to predict the number of photons 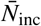 at photon energy *E*_inc_ required per pixel when attempting to detect a mass concentration 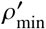 of any specified element.

### 3.2. Corresponding radiation dose to a matrix material

In many studies, one is measuring a low mass concentration 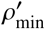 of a specified element present in a higher-concentration matrix material; one example involves study of the role of zinc in oocytes and embryos during fertilization [41–43]. High radiation doses can lead to morphological changes and mass loss in the organic materials in cells and tissues [44], so it is also important to provide an estimate of the radiation dose imparted to a matrix material (which we denote with the subscript “mat”) associated with irradiation with 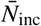 photons per area. To do so, we first consider the incident fluence 𝔉_inc_ of

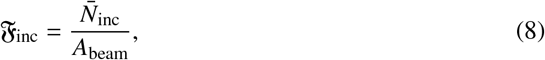

where *A*_beam_ is the area of the incident beam (the beam spot size). Inserting the result of Eq. 7 into this expression yields

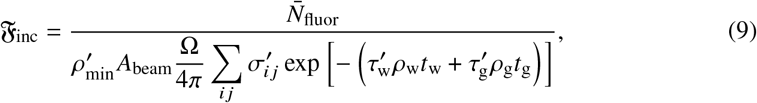

where 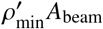 is also known as the minimum detectable mass [20]. Skin dose *D*_skin_ is the radiation dose delivered to the beam-facing surface of a matrix material; it can be found from [20, 44]

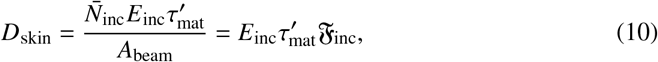

where [44]

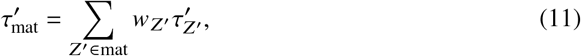

and where the final form of Eq. 10 comes from Eq. 8. In the above two equations, 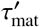 is the total mass photoionization cross section of the sample matrix, and *w*_*Z*_′ is the weighting coefficient accounting for the atom number fraction of each element *Z*^′^ present in the matrix material (that is, for each *Z*^′^ ∈ mat). If the sample is tilted by angle θ relative to the transverse of the incident beam (so as to balance between XRF self-absorption minimization and beam broadening), then the beam width as seen by the tilted sample pixels along one direction increases by a factor of cos θ. In this case, Eq. 10 becomes

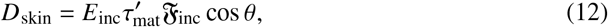

leading to a factor cos θ drop in the skin dose. Substituting Eq. 9 into the above equation results in

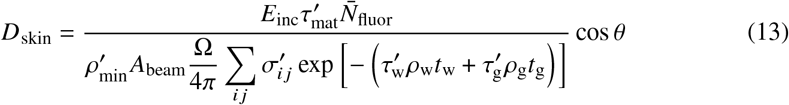

as the skin dose.

It is common in x-ray imaging calculations to represent biological specimens as being comprised of a model protein with the compositional average of all 20 amino acids, with a stoichiometric composition of H_48.6_C_32.9_N_8.9_O_8.9_S_0.6_ [45] (the density does not need to be specified; see Sec. S2 in Supplement 1). We used that protein as the matrix material in the skin dose calculations described in Sec. 3.3.

### 3.3. Numerical example: low-concentration elements in a protein matrix

In Sec. 3.1, we derived the minimum number 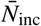 of incident x-ray photons per pixel (Eq. 7) required for detection of an elemental concentration 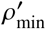. As noted in Sec. 3.1, those derivations were performed under the assumption that the detection of 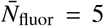 photons per pixel is sufficient for elemental detection (this point is discussed further in Sec. 3.4). Because this covers most SXFM studies today, our calculations only include x-ray fluorescence from *K* and *L* subshells; our approach could be extended to *M* subshells and beyond if desired. From the calculations of 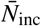, we also computed the corresponding matrix material skin dose *D*_skin_ (Eq. 13) as described in Sec. 3.2. We used those results to obtain numerical estimates representative of typical experiments while assuming the following:

- The specimen matrix is the model protein of stoichiometric composition H_48.6_C_32.9_N_8.9_O_8.9_S_0.6_ as discussed in Sec. 3.2.
- The specimen is illuminated with a single incident photon energy *E*_inc_.
- The specimen contains trace elements *Z* between _10_Ne and _92_U, all at an areal mass concentration of 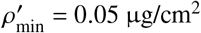. This value of 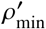 is representative of the limit of detection in an SXFM experiment.
- The specimen is housed in a vacuum environment so that *t*_g_ = 0.
- For each element *Z*, a fluorescence signal with 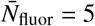 photons per pixel must be counted by a windowless detector with an acceptance solid angle of Ω = 1.35 sr.

These assumptions were sufficient to calculate the required number of incident photons 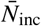 per pixel. For computing the resulting skin dose *D*_skin_ in the matrix material, we used a value of *A*_beam_ corresponding to a circular beam focus 40 nm in diameter, and we assumed that the specimen was at normal incidence to the beam so that θ = 0.

Our calculations utilized tabulations provided by the *xraylib* fundamental parameter database [30]. That database contains information relevant for *K, L*, and *M* shell fluorescence and involves a complete XRF forward model that accounts for the excitation dependence of mass photoionization partial cross sections (PCSes), CK transitions, and cascade effects. The *xraylib* database does not include some weak fluorescence lines, like *K*α_3_ and *L* β_2_, that are formally forbidden by selection rules in single electron theory [46] (though they can in fact be weakly present). The database also does not include non-radiative transitions such as super Coster-Kronig (SCK) transitions. These omitted parameters would not lead to noticeable changes in our results if they were somehow to be included.

With the above assumptions and fundamental parameter tabulations in hand, we show in Fig. 1 our calculated values of 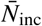 (a) and *D*_skin_ (b) as a function of atomic number *Z*, as well as a function of individual incident photon energies *E*_inc_ over the range of 4 to 34 keV. The values of 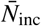 and *D*_skin_ are shown using a false color map, with the color map scale shown at right. One can think of this as a topographical map of terrain, with contour lines at altitude intervals. The contour lines are labeled with numbers *C*, which correspond to values of 10^*C*^ for 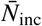 and *D*_skin_. The calculations utilized *K* and *L* fluorescence lines only, so the white region at lower right reflected incident photon energies *E*_inc_ that were too low to reach the threshold for exciting *L* line fluorescence. In a similar fashion, the plots showed a “topographical cliff”, or sharp decrease, in both 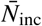 (a) and *D*_skin_ (b) when *E*_inc_ increased to reach the threshold for exciting *K* fluorescence; this cliff started at (*Z* = 20, *E*_inc_ = 4 keV) and rose to (*Z* = 54, *E*_inc_ = 34 keV).

**Fig. 1.**
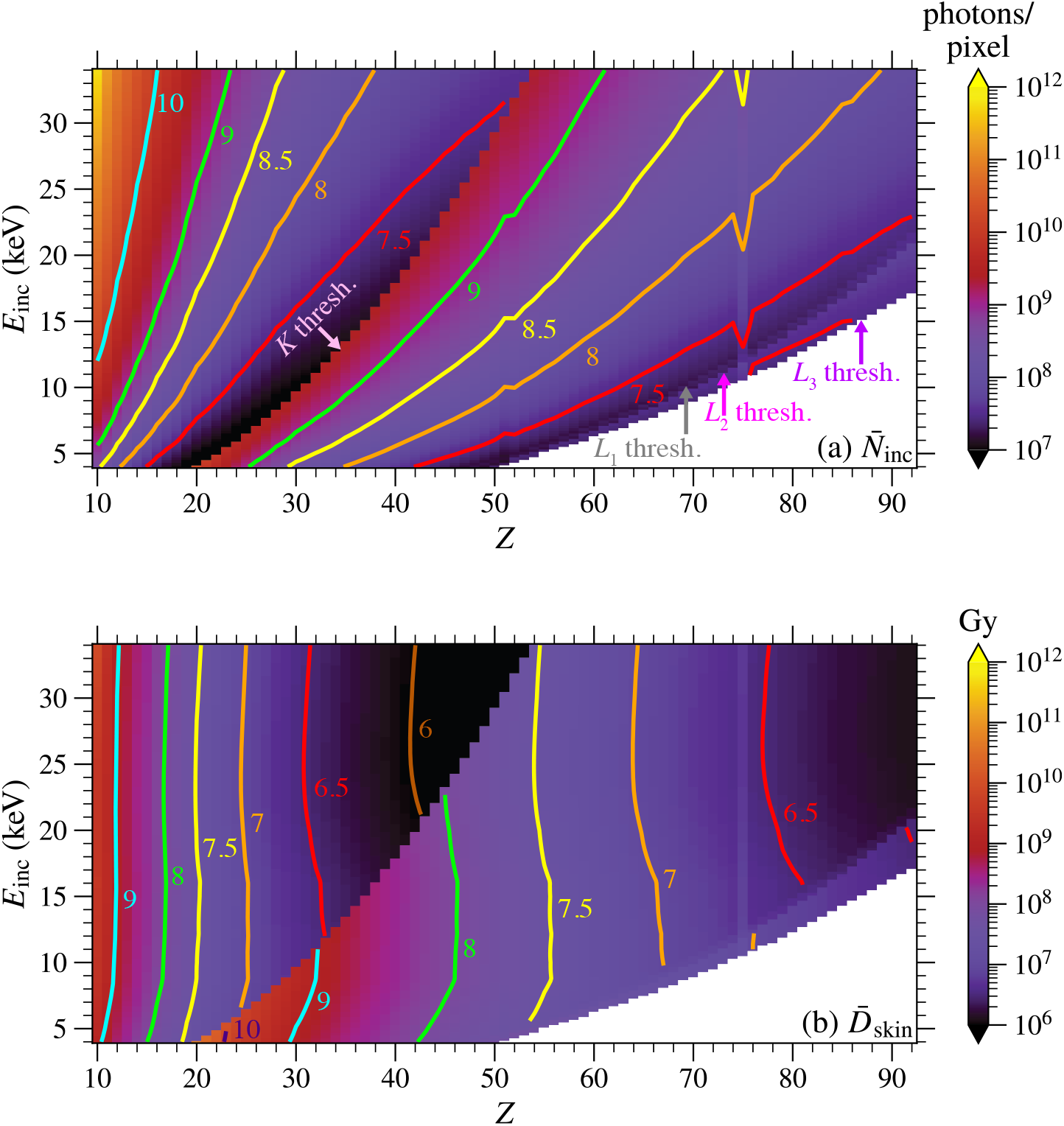
Combined false color maps and contour plots of the expected minimum number of incident photons 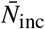 per pixel (a) and skin dose *D*_skin_ imparted (b) for element detection in vacuum. These values are shown versus trace element atomic number *Z* and individual incident photon energies *E*_inc_. These calculations were carried out for a limit of detection of 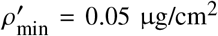 and for the detection of 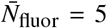 x-ray photons per pixel summed over all accessible *K* and *L* fluorescence emission lines. We assumed the x-ray fluorescence detector was windowless and had a solid angle of collection of Ω = 1.35 sr. The skin dose *D*_skin_ (b) associated with 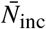 was calculated assuming a model protein composition of H_48.6_C_32.9_N_8.9_O_8.9_S_0.6_ [45] and a focused beam diameter of 40 nm. These calculations included all the effects related to photoionization partial cross sections as described in (Sec. 3.1). As *Z* increased, the contributions to 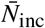 and *D*_skin_ for a particular subshell abruptly changed when *E*_inc_ hit and exceeded that subshell”s absorption edge (labeled as “thresh.”). The white regions in each panel exist due to an individual value of *E*_inc_ not being high enough to excite the *L*_3_ subshell, as well as due to the exclusion of XRF events stemming from shells greater than *L*_3_ from our calculations. This calculation employed tabulated data from *xraylib* [30].

### 3.4. Experimental validation

The calculations of the minimum number of incident photons per pixel 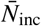 of Eq. 7 assumed zero background when detecting 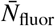 fluorescence photons per pixel. There are relatively few published reports that provide sufficient detail to make additional experimental comparisons between the number of incident photons 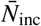 per pixel and a minimum detected areal mass concentration 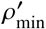. There is sufficient information, however, in one recent experiment [47] of *K* fluorescence of five low-concentration elements present in a scanning fluorescence x-ray microscopy (SFXM) experiment.

The experiment involved a 10 µm thick section of dehydrated mouse kidney tissue mounted on a Si_3_N_4_ window, imaged at beamline 8-BM-B at the Advanced Photon Source at Argonne National Lab. In a typical scan with a per-pixel imaging time of *t*_dwell_ = 50 ms, the sample was illuminated with 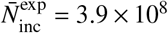 photons per pixel (±5%) at *E*_inc_ = 10 keV photon energy. The full fluorescence spectrum was obtained using a 7-element energy dispersive detector with an acceptance solid angle of Ω = 1.35 sr and an entrance window of *t*_w_ = 25 µm thick beryllium. The values of Ω and *E*_inc_ here correspond to the calculation assumptions described in Sec. 3.3. The recorded spectrum was analyzed with the M-BLANK software package [27] using spectral data obtained from a sample-free Si_3_N_4_ window for background subtraction, as well as elemental areal mass concentrations obtained by comparison with fluorescence signals obtained from an AXO 10X thin film standard (RF8-200-S2454, Applied X-ray Optics, GmbH). In that experiment, the elements _15_P, _16_S, _20_Ca, _26_Fe, and _28_Ni were all present at a wide range of concentrations. The forward model of Eq. 1 assumed no background other than from Poisson fluctuations stemming from the detection of fluorescence photons themselves; this well-approximated the case of the selected elements since their true XRF signals were either originally much stronger than that of the experimental background or enough of the background was subtracted out when initially fitting raw fluorescence spectra.

For the five selected elements, we defined the limit of detection 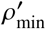 via background–corrected fits to the fluorescence of the Si_3_N_4_ window. After measuring the fluorescence emitted from an empty Si_3_N_4_ window, we averaged the resulting spectrum over all pixels to acquire a representative background spectrum. This average background was then subtracted at every pixel of the kidney section scan, and we fit the acquired difference spectra using the same M-BLANK parameterized peak model employed for the sample data [48]. This yielded a population of fitted, background-corrected signals (expressed as calibrated areal mass concentrations *ρ*^′^) across all substrate pixels for each element. We took the standard deviations σ_*ρ*_′ of those distributions as the noise levels, and defined the limit of detection (LOD) to be 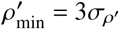. This approach provided an element-specific, data-driven estimate of 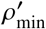 under the same experimental conditions and fitting model as the sample measurements.

To obtain measures of the total number of fluorescence photons *N*_fluor_ collected at each pixel within 1% of the per-element 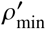 determined above, we defined energy windows for summing *K*α and *K* β photons using *xraylib*-tabulated line energies *E*_*i j*_ [30] combined with an empirically calibrated detector response function. We specified the nominal photon energies of the *K*α_1_, *K*α_2_, *K* β_1_, and (where relevant) *K* β_2_ lines and treated those values as line centroids. Afterward, for each detector element, we parameterized the line energy resolution Δ*E*_*i j*_ as an energy-dependent full width at half maximum (FWHM) via a standard Fano-limited model of [49]

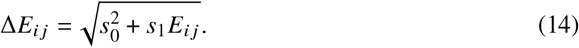

In the above equation, *s*_0_ and *s*_1_ are detector element-specific parameters obtained by fitting the measured detector element response to multiple fluorescence lines in the same dataset. We then defined the integration window bounds 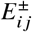 each fluorescence line *i j* according to

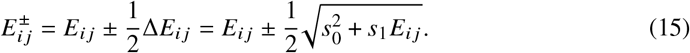

Because all values of *s*_0_ and *s*_1_ were extracted directly from Si_3_N_4_ scans rather than from a fixed lookup table, the resulting energy windows accurately reflected the actual detector performance under the specific beamline and low-count-rate conditions used in the experiment. For calcium, where the *K*α peaks have some overlap with potassium *K* β lines when the peak broadening of the energy-dispersive detector is taken into account, we mitigated interference by excluding pixels in which the fitted potassium concentration map exceeded its own value of 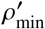 scaled up by the expected XRF intensity ratio *F*_*K* α_ / *F*_*K*β_ = 8.65 [30]. This ensured that the calcium photon statistics near the calcium limit of detection were not dominated by potassium *K* β spill-over [50].

After calculating all energy windows, we summed up the total number of photons collected over all detector elements for each pixel within those windows to obtain an aggregate number *N*_fluor_ of XRF photons collected over all detector elements for each pixel around 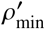. Ultimately, all of this resulted in a distribution of collected fluorescence photons *N*_fluor_ across all selected pixels, shown as histograms in Fig. 2. From these histograms of probability densities for each element, we obtained the mean number of XRF photons 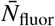collected per pixel.

**Fig. 2.**
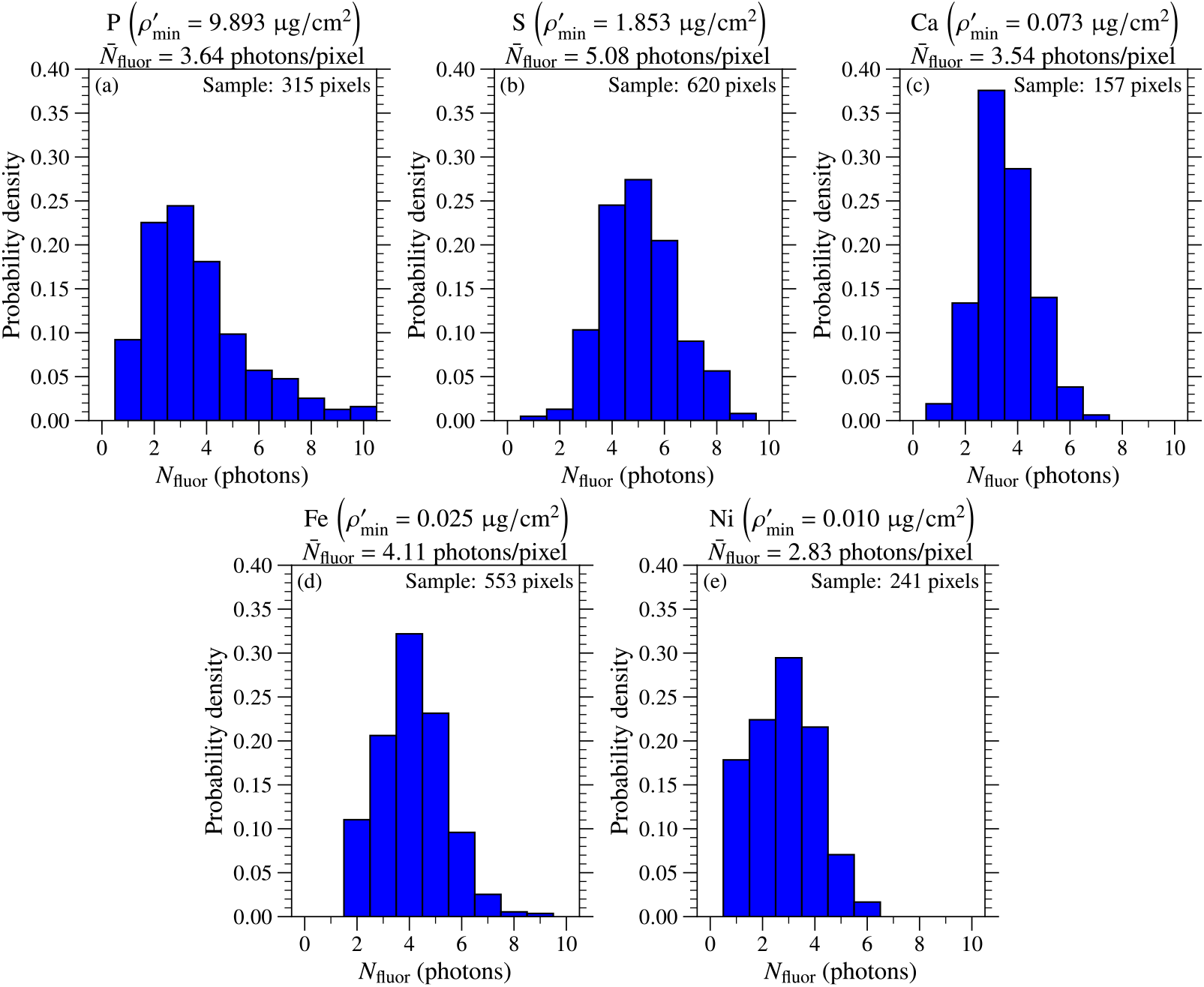
The distribution of total x-ray fluorescence photons detected in each pixel around different elemental limits of detection (LODs) 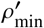in our experiment at beamline 8-BM-B [47]. Shown here are histograms of probability densities for several elements with respect to the total number *N*_fluor_ of XRF photons detected for sample pixels within 1% of each element”s LOD, which we obtained by summing up the contributions from all detector elements. From those distributions, we were able to calculate the mean minimum number of x-ray fluorescence photons 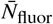 detected per pixel.

In Table 2, we show for the five selected elements both the limits of detection 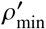 and the mean number of detected x-ray fluorescence photons 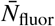 obtained from experimental data using the procedures described above. This table also shows 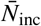 as calculated from Eq. 7 using those experimental values of 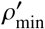 and 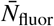 when neglecting any attenuation due to gas in the sample environment. Since the specimen was illuminated with 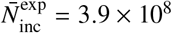 photons perpixel, we also show the ratio 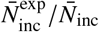 for when both excluding and including air attenuation effects. The ratio 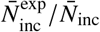 for *t*_g_ = 0 showed especially large discrepancies for the lower-*Z* elements with XRF emission at lower photon energies. Much of that was attributed to the fact that the capillary optic experimental apparatus described above did not include a helium gas enclosure around the specimen, leading to an air gap *t*_g_ between the sample and the entrance window of the energy-dispersive detector. While we did not have an exact measurement of the air gap, we estimated it to be about *t*_g_ = 5 cm. Accounting for that gap brought the ratio 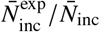 closer to a factor of 5–12 for those elements.

**Table 2.**
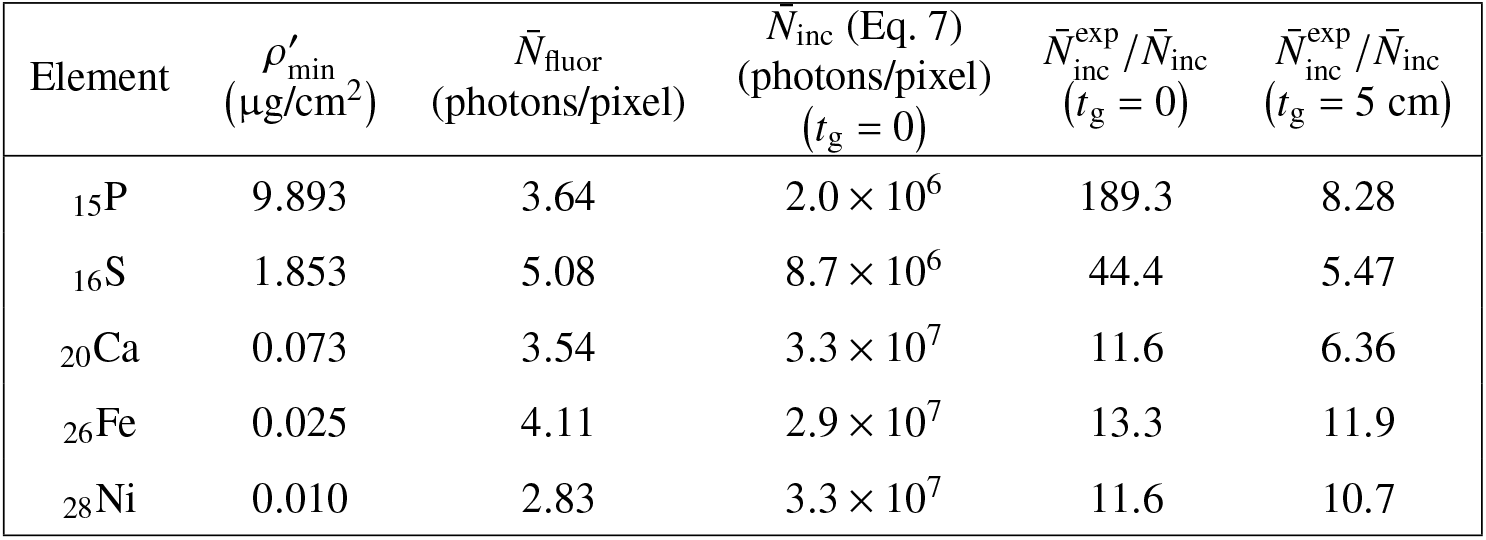
Comparison of experimental (with 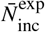 from Sec. 3.4) and calculated values for mass concentrations 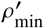 and detected number of fluorescent photons 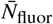, along with the number of incident photons 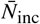. The 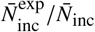 omparison was made both in the absence of an absorbing gas path in air (*t*_g_ = 0) and with the estimated value of *t*_g_ = 5 cm

## 4. Effects of subshell excitation models

The calculation results shown in Table 2 incorporated the excitation dependence of subshell mass photoionization cross sections 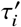, the existence of CK transitions, and the existence of both radiative and nonradiative cascade effects (see Secs. 1 and 2). We now consider the changes that would arise with less-exact calculations. For these illustrations, we used a single incident photon energy of *E*_inc_ = 10 keV, and we assumed that 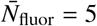 fluorescent photons per pixel were required to detect an areal concentration of 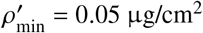 when using a windowless EDS detector.

To illustrate the shortcomings of using the simpler “jump ratio” model of Eq. 4, we show in Fig. 3 differences in requirements for the minimum number of excitation photons 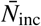 with and without this simpler model. As can be seen, the “jump ratio” approximation lead to only small differences in 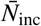 when detecting *K* fluorescence lines, but it lead to erroneously high estimates of 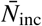 for the *L* shell. This was especially true when relying on *L*_1_ line emission for elemental detection, as CK transitions cannot occur in that subshell. For that case, the error increased with greater differences between the incident photon energy *E*_inc_ and the absorption edge energy 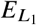 of a particular element. Differences like these have been experimentally observed [36, 38]. In one example, inaccurate quantification of the thickness of a palladium (Pd) thin film was observed as the incident photon energy *E*_inc_ was increased well beyond the energy of each of the three *L* absorption edges; this demonstrated the inaccuracy of the jump ratio approach, in particular when using *L*_1_ fluorescence emission lines [37].

**Fig. 3.**
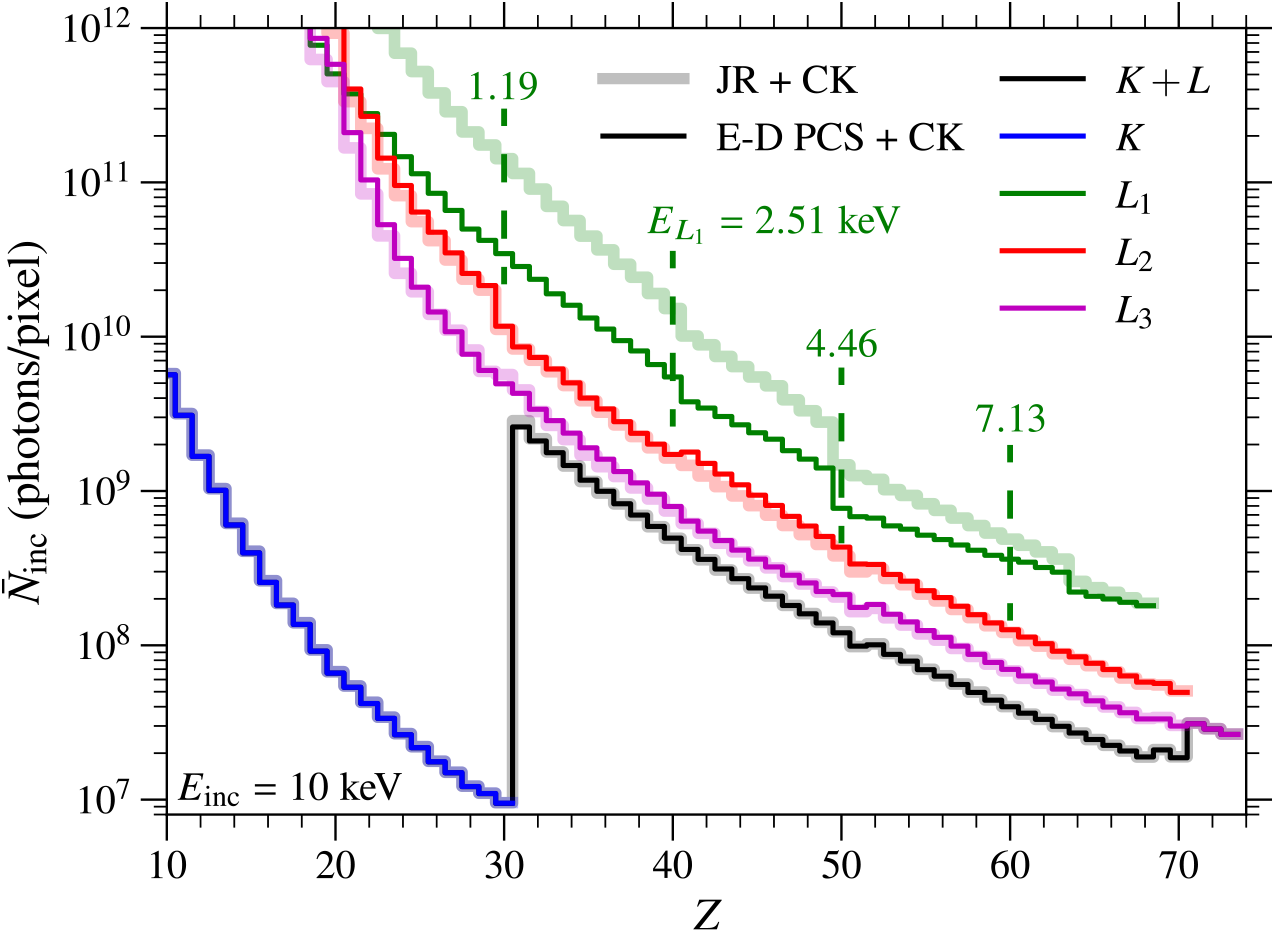
Subshell-specific calculations of the minimum number of incident photons 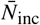 per pixel for a fixed incident photon energy of *E*_inc_ = 10 keV, with and without the “jump ratio” approximation. The (E-D PCS + CK) calculations were carried out using excitation-dependent mass photoionization partial cross sections, while the less-accurate (JR + CK) calculations were carried out using the “jump ratio” approximation of Eq. 4. In both cases, Coster-Kronig transitions were included (CK). As can be seen, the “jump ratio” approximation leads to inaccurately high calculated values of 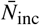 in particular when considering fluorescence from *L*_1_ lines when the excitation energy *E*_inc_ is well above the edge energy 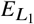. We assumed a windowless EDS detector and a vacuum environment for these calculations so as to not affect 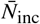 at lower *L* fluorescence emission energies. Tabulated data from *xraylib* [30].

The exclusion of cascade effects can also lead to erroneous estimates of the required number of incident photons 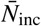 when considering *L* fluorescence emission lines, as shown in Fig. 4. For that comparison, 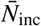 was lower for all values up until _30_Zn, at which point *E*_inc_ = 10 keV is too low to excite the *K* edges of higher *Z* elements. Because SCK transitions in the *L* shell were omitted, all values of 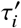 above Zn were the same as their non-cascading counterparts. (Again, those values would not change significantly if they were included somehow.)

**Fig. 4.**
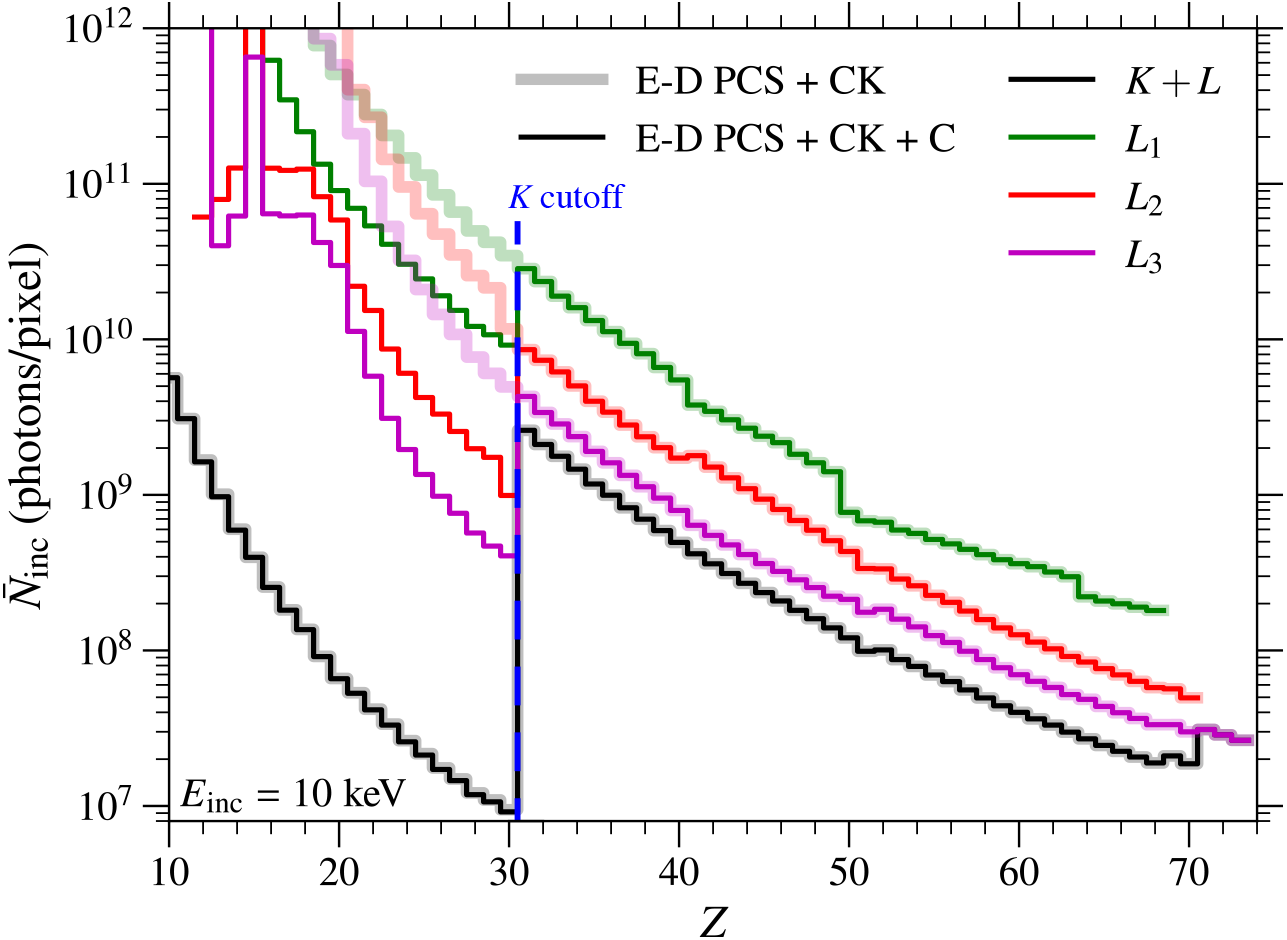
Subshell-specific calculations of the minimum number of incident photons 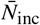 per pixel for a fixed incident photon energy of *E*_inc_ = 10 keV with and without the incorporation of cascade effects. Cascading due to *K* shell photoionization ceased when the *K* edge energy (labeled “*K* cutoff”) exceeded *E*_inc_, which occurs for *Z* ≥ 31 when *E*_inc_ = 10 keV. Double vacancies caused by electrons emitted during *L* shell super Coster-Kronig (SCK) transitions were omitted; thus, there were no cascade effects at all past the “*K* cutoff” shown. Shown here are the results when cascade effects (both radiative and nonradiative) are both considered (E-D PCS + CK + C), and ignored (E-D PCS + CK). In both cases, we used energy-dependent partial cross sections (E-D PCS) and included Coster-Kronig transitions (CK). We assumed a windowless EDS detector and a vacuum environment for these calculations so as to not affect 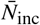 at lower *L* fluorescence emission energies. Tabulated data from *xraylib* [30].

In Figs. 3 and 4, we observed differences in the estimates of 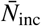 based only on considering *L* subshells. However, those differences effectively disappeared when considering the sum of all *K* and *L* shell fluorescence contributions.

## 5. Discussion

The experimental comparison of Sec. 3.4 highlighted how our theoretical predictions relate to practical detection limits. This showed that the calculations, although optimistic, were not disconnected from reality. In experimental results, the observed limits of detection 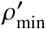 decreased with increasing atomic number *Z*, matching the expected trend: as the incident photon energy *E*_inc_ reaches and exceeds an element”s absorption edge, the total mass photoionization cross section τ^′^ for that element increases, making photoionization (and therefore x-ray fluorescence) more probable. Correspondingly, the empirical XRF histograms of Fig. 2 showed that detecting roughly three to six fluorescence photons per pixel was consistently sufficient to reach the operational limits of detection across all elements examined. Together, these observations showed that the theoretical false color map/contour plot predictions of Fig. 1 yielded estimates of 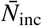 approximately consistent with observations in one experiment, though the experimental values were 5–12 times higher.

## 6. Conclusion

We have presented here an approach to estimate the minimum number of incident photons 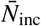 per pixel required for the detection of low-concentration elements when using a single incident photon energy *E*_inc_ to excite x-ray fluorescence from many different elements *Z*, which is representative of most experiments in scanning fluorescence x-ray microscopy (SFXM). Earlier calculations [22] assumed the use of an incident photon energy just above an element”s absorption edge, ideal for detecting just one element. In addition, we made use of *xraylib* [30], which provides computer-accessible tabulations of all relevant parameters and thus allows for more complete calculations. As a result, our model accounted for the incident energy dependence of mass photoionization partial cross sections, Coster-Kronig transitions, cascade effects, and attenuation due to detector windows and gas within a sample environment. As technology upgrades at synchrotrons lead to higher photon brightness, estimates of the limits of detection play an increasingly important role in the planning of SXFM experiments, laying the groundwork for next generation discoveries in many areas including inorganic physiology.

## Acknowledgment

We thank Philipp Hönicke for discussions that led us to appreciate the factors that come into play when using excitation energies *E*_inc_ that are well above the absorption edges of particular elements, especially when *L* fluorescence lines are considered.

## Data Availability Statement

Experimental data used in this manuscript can be found in our previous work [47]. The code we developed for our calculations can be found on Github: https://github.com/bwr0835/xray_fluor_contrast.

Note: Contour values correspond to base-10 exponents.

## References

1. C. G. Barkla, “The spectra of the fluorescent Röntgen radiations,” Philos. Mag. 22, 396–412 (1911).

2. H. G. J. Moseley, “The high-frequency spectra of the elements,” Philos. Mag. 26, 1024–1034 (1913).

3. H. G. J. Moseley, “The high-frequency spectra of the elements. Part II,” Philos. Mag. 27, 703–713 (1914).

4. P. Horowitz and J. A. Howell, “A scanning x-ray microscope using synchrotron radiation,” Science 178, 608–611 (1972).

5. C. J. Sparks, Jr., “X-ray fluorescence microprobe for chemical analysis,” in Synchrotron Radiation Research, H. Winick and S. Doniach, eds. (Plenum Press, New York, 1980), chap. 14, pp. 459–512.

6. K. W. Jones, B. M. Gordon, A. L. Hanson, et al., “Application of synchrotron radiation to elemental analysis,” Nucl. Instruments Methods Phys. Res. B 3, 225–231 (1984).

7. T. Paunesku, S. Vogt, J. Maser, et al., “X-ray fluorescence microprobe imaging in biology and medicine,” J. Cell. Biochem. 99, 1489–1502 (2006).

8. C. Fahrni, “Biological applications of x-ray fluorescence microscopy: exploring the subcellular topography and speciation of transition metals,” Curr. Opin. Chem. Biol. 11, 121–127 (2007).

9. M. J. Pushie, I. J. Pickering, M. Korbas, et al., “Elemental and chemically specific x-ray fluorescence imaging of biological systems,” Chem. Rev. 114, 8499–8541 (2014).

10. M. D. de Jonge and S. Vogt, “Hard x-ray fluorescence tomography -an emerging tool for structural visualization,” Curr. Opin. Struct. Biol. 20, 606–614 (2010).

11. D. Sayre, J. Kirz, R. Feder, et al., “Transmission microscropy of unmodified biological materials: comparative radiation dosages with electrons and ultrasoft x-ray photons,” Ultramicroscopy 2, 337–341 (1977).

12. D. Rudolph, G. A. Schmahl, and B. Niemann, “Amplitude and phase contrast in x-ray microscopy,” in Modern Microscopies, P. J. Duke and A. G. Michette, eds. (Plenum, New York, 1990), pp. 59–67.

13. G. Schneider, “Cryo x-ray microscopy with high spatial resolution in amplitude and phase contrast,” Ultramicroscopy 75, 85–104 (1998).

14. M. R. Howells, T. Beetz, H. N. Chapman, et al., “An assessment of the resolution limitation due to radiation-damage in x-ray diffraction microscopy,” J. Electron Spectrosc. Relat. Phenom. 170, 4–12 (2009).

15. M. Du and C. Jacobsen, “Relative merits and limiting factors for x-ray and electron microscopy of thick, hydrated organic materials,” Ultramicroscopy 184, 293–309 (2018).

16. Q. Shen, I. Bazarov, and P. Thibault, “Diffractive imaging of nonperiodic materials with future coherent x-ray sources,” J. Synchrotron Radiat. 11, 432–438 (2004).

17. A. Schropp and C. G. Schroer, “Dose requirements for resolving a given feature in an object by coherent x-ray diffraction imaging,” New J. Phys. 12, 035016 (2010).

18. B. De Samber, M. J. Niemiec, B. Laforce, et al., “Probing intracellular element concentration changes during neutrophil extracellular trap formation using synchrotron radiation based x-ray fluorescence,” PLoS ONE 11, e0165604 (2016).

19. F. Adams, B. Vekemans, G. Silversmit, et al., “Microscopic x-ray fluorescence analysis with synchrotron radiation sources,” (Springer, 2011), Handbook of Nuclear Chemistry, pp. 1737–1759.

20. J. Kirz, D. Sayre, and J. Dilger, “Comparative analysis of x-ray emission microscopies for biological specimens,” Ann. New York Acad. Sci. 306, 291–305 (1978).

21. J. Kirz, “Specimen damage considerations in biological microprobe analysis,” in Scanning Electron Microscopy, vol. 2 (SEM Inc., Chicago, 1980), pp. 239–249.

22. J. Kirz, “Mapping the distribution of particular atomic species,” Ann. New York Acad. Sci. 342, 273–287 (1980).

23. D. Z. Zee, K. W. MacRenaris, and T. V. O’Halloran, “Quantitative imaging approaches to understanding biological processing of metal ions,” Curr. Opin. Chem. Biol. 69, 102152 (2022).

24. S. Vogt, “MAPS: a set of software tools for analysis and visualization of 3D x-ray fluorescence data sets,” J. de Physique IV 104, 635–638 (2003).

25. V.A. Solé, E. Papillon, M. Cotte, et al., “A multiplatform code for the analysis of energy-dispersive x-ray fluorescence spectra,” Spectrochimica Acta Part B: At. Spectrosc. 62, 63–68 (2007).

26. C. G. Ryan, R. Kirkham, R. M. Hough, et al., “Elemental x-ray imaging using the Maia detector array: The benefits and challenges of large solid-angle,” Nucl. Instruments Methods Phys. Res. A 619, 37–43 (2010).

27. A. M. Crawford, A. Deb, and J. E. Penner-Hahn, “M-BLANK: a program for the fitting of x-ray fluorescence spectra,” J. Synchrotron Radiat. 26, 497–503 (2019).

28. R. A. Van Grieken and A. A. Markowicz, Handbook of X-ray Spectrometry, vol. 29 of Practical Spectroscopy (Marcel Dekker, New York, 2002), 2nd ed.

29. E. De Pauw, P. Tack, and L. Vincze, “A review of laboratory, commercially available, and facility based wavelength dispersive x-ray fluorescence spectrometers,” J. Anal. At. Spectrom. 39, 310–329 (2024).

30. T. Schoonjans, A. Brunetti, B. Golosio, et al., “The xraylib library for X-ray-matter interactions. Recent developments,” Spectrochimica Acta Part B: At. Spectrosc. 66, 776–784 (2011).

31. J. Sherman, “The theoretical derivation of fluorescent x-ray intensities from mixtures,” Spectrochimica Acta 7, 283–306 (1955).

32. A. C. Thompson, D. T. Attwood, E. Gullikson, et al., X-ray Data Booklet (Lawrence Berkeley National Laboratory, University of California, Berkeley, CA, 2009), rev. 3 ed.

33. L. H. Martin, “The efficiency of K series emission by K ionised atoms,” Proc. Royal Soc. A 115, 420–442 (1927).

34. A. H. Compton and S. K. Allison, X-Rays in Theory and Experiment (D. Van Nostrand, New York, 1935), 2nd ed.

35. J. H. Scofield, “Theoretical photoionization cross sections from 1 to 1500 keV,” Tech. Rep. UCRL-51326, University of California (1973).

36. P. Hönicke, M. Kolbe, M. Müller, et al., “Experimental verification of the individual energy dependencies of the partial ??-shell photoionization cross sections of Pd and Mo,” Phys. Rev. Lett. 113, 163001 (2014).

37. P. Hönicke, M. Kolbe, and B. Beckhoff, “What are the correct L-subshell photoionization cross sections for x-ray spectroscopy?” X-ray Spectrom. 45, 207 (2016).

38. P. Hönicke, “A novel and holistic approach for experimental x-ray fundamental parameter determination–the Ru L-shell,” New J. Phys. 25, 073012 (2023).

39. W. Bambynek, B. Crasemann, R. W. Fink, et al., “X-ray fluorescence yields, Auger, and Coster–Kronig transition probabilities,” Rev. Mod. Phys. 44, 716–813 (1972).

40. L. A. Currie, “Limits for qualitative detection and quantitative determination. Application to radiochemistry,” Anal. Chem. 40, 586–593 (1968).

41. E. L. Que, R. Bleher, F. E. Duncan, et al., “Quantitative mapping of zinc fluxes in the mammalian egg reveals the origin of fertilization-induced zinc sparks,” Nat. Chem. 7, 130 – 139 (2015).

42. B. Y. Kong, F. E. Duncan, E. L. Que, et al., “The inorganic anatomy of the mammalian preimplantation embryo and the requirement of zinc during the first mitotic divisions,” Dev. Dyn. 244, 935–947 (2015).

43. J. L. Balough, T. V. O’Halloran, F. E. Duncan, and T. K. Woodruff, “Inorganic profiles of preimplantation embryos reveal a role for zinc in blastocyst development,” bioRxiv p. 2025.05.14.654061 (2025).

44. C. Jacobsen, X-ray Microscopy (Cambridge University Press, Cambridge, UK, 2020).

45. R. A. London, M. D. Rosen, and J. E. Trebes, “Wavelength choice for soft x-ray laser holography of biological samples,” Appl. Opt. 28, 3397–3404 (1989).

46. N. A. Dyson, X-rays in Atomic and Nuclear Physics (Cambridge University Press, Cambridge, UK, 1973), 2nd ed.

47. B. Roter, A. M. Crawford, Q. Jin, et al., “Multifunctional bending magnet beamline with a capillary optic for x-ray fluorescence studies of metals in tissue sections,” bioRxiv (2025).

48. A. M. Crawford, “X-ray fluorescence data analysis using a blank correction approach,” in Advances in X-ray Analysis, vol. 63 L. Brehm and M. Schmeling, eds. (JCPDS-International Centre for Diffraction Data, 2020), pp. 180–193.

49. D. M. Schlosser, P. Lechner, G. Lutz, et al., “Expanding the detection efficiency of silicon drift detectors,” Nucl. Instruments Methods Phys. Res. A 624, 270–276 (2010).

50. A. M. Crawford, N. J. Sylvain, H. Hou, et al., “A comparison of parametric and integrative approaches for x-ray fluorescence analysis applied to a Stroke model,” J. Synchrotron Radiat. 25, 1780–1789 (2018).

