## Supplementary material for "X-ray fluorescence microscopy exposure estimates using a single excitation energy"

### 1. Clarification on x-ray nomenclature

For simplicity in mathematical notation, in the main text, we refer to fluorescence lines by the indices $ij$, where $i$ and $j$ are electron vacancy states involved in x-ray fluorescence transitions. When photoelectric absorption leads to an initial vacancy in the $i = 1s$ state and an electron drops down so that the $j = 2p_{3/2}$ state represents the subsequent vacancy, the energy difference is released via an x-ray fluorescence photon. This fluorescence event can be described using either the IUPAC notation [1] of $K$-$L_3$ or the equivalent Siegbahn notation [2] of $K\alpha_1$. Additionally, when we refer to $K$ or $L_1$ shell emission, $K$ or $L_1$ fluorescence, or fluorescence emitted from $K$ or $L_1$, we are describing x-ray fluorescence events due to vacancies being filled in the $K$ or $L_1$ shell, respectively. This is less specific than referring to a particular fluorescence line like $K\alpha_1$.

### 2. Total mass absorption coefficient for mixtures

In Sec. 3.2 of the main manuscript document, we defined the radiation skin dose $D_{\text{skin}}$ in Eqs. 10 and 11, which (when ignoring scattering) is proportional to a sample matrix's total mass photoionization cross section $\tau'_{\text{mat}}$. Another approach for calculating skin dose involves the density $\rho_{\text{mat}}$ and linear absorption coefficient $\mu_{\text{mat}}$ of the matrix material, leading to an expression of [3, 4]

$$D_{\text{skin}} = \frac{\bar{N}_{\text{inc}} E_{\text{inc}} \mu_{\text{mat}}}{\rho_{\text{mat}} A_{\text{beam}}} = \frac{E_{\text{inc}} \mu_{\text{mat}} \mathfrak{F}_{\text{inc}}}{\rho_{\text{mat}}}. \tag{S1}$$

However, the linear absorption coefficient can be written as

$$\mu_{\text{mat}} = \frac{N_A \rho_{\text{mat}}}{A_{\text{mat}}} \tau_{\text{mat}}, \tag{S2}$$

where

$$\tau_{\text{mat}} = 2 r_e \frac{hc}{E_{\text{inc}}} f_{2,\text{mat}}. \tag{S3}$$

In the above two equations, $\tau_{\text{mat}}$ and $A_{\text{mat}}$ are the total atomic photoionization cross section and molar mass of the matrix, respectively, $r_e$ is the classical electron radius, $N_A$ is Avogadro's number, $h$ is Planck's constant, $c$ is the speed of light in vacuum, and $f_{2,\text{mat}}$ is the total imaginary part of the complex number of oscillator modes the atoms of the matrix [4, 5]. Because $\mu_{\text{mat}}$ is dependent on $\rho_{\text{mat}}$, the matrix material's density does not actually factor into the calculated dose. Mass attenuation coefficients and cross sections are readily available in tabulations and are independent of $\rho_{\text{mat}}$.

The linear absorption coefficients $\mu_{Z'}$ for each element in the matrix can be written in terms of their corresponding mass attenuation coefficients $\mu'_{Z'}$ via

$$\mu_{Z'} = \mu'_{Z'} \rho_{Z'}, \tag{S4}$$

where

$$\mu'_{Z'} = \left(\frac{\mu}{\rho}\right)_{Z'} = \frac{N_A}{A_{Z'}} \sigma_{Z'}. \tag{S5}$$

In Eq. S5, $A_{Z'}$ and $\sigma_{Z'}$ are the molar mass and total atomic cross section for atomic number $Z'$, respectively. [Notationally, $(\mu/\rho)_{Z'} \equiv \mu_{Z'}/\rho_{Z'}$.] Uniform sample matrices allow for the calculation of $\rho_{Z'}$ as [4]

$$\rho_{Z'} = \frac{s_{Z'} A_{Z'}}{A_{\text{mat}}} \rho_{\text{mat}} = w_{Z'} \rho_{\text{mat}}, \tag{S6}$$

where

$$w_{Z'} = \frac{s_{Z'} A_{Z'}}{A_{\text{mat}}}. \tag{S7}$$

Here, $s_{Z'}$ and $w_{Z'}$ are the stoichiometric and mass fractional weighting coefficients of atomic number $Z'$, respectively. One can obtain $\mu_{\text{mat}}$ by inserting Eq. S6 into Eq. S4, and summing over all $Z' \in \text{mat}$. All of this yields

$$\mu_{\text{mat}} = \sum_{Z' \in \text{mat}} \mu_{Z'} = \rho_{\text{mat}} \sum_{Z' \in \text{mat}} w_{Z'} \mu'_{Z'}. \tag{S8}$$

Dividing the above equation by $\rho_{\text{mat}}$ gives

$$\mu'_{\text{mat}} \equiv \frac{\mu_{\text{mat}}}{\rho_{\text{mat}}} = \sum_{Z' \in \text{mat}} w_{Z'} \mu'_{Z'}. \tag{S9}$$

When neglecting scattering, $\sigma_{Z'} = \tau_{Z'}$, and $\mu'_{Z'} = (N_A / A_{Z'}) \tau_{Z'} = \tau'_{Z'}$, with $\tau_{Z'}$ being the total atomic photoionization cross section for atomic number $Z'$. Therefore,

$$\tau'_{\text{mat}} = \sum_{Z' \in \text{mat}} w_{Z'} \tau'_{Z'}. \tag{S10}$$

This complements the mixture rule [4, 6, 7] applied to linear absorption coefficients $\mu_{\text{mat}}$.

### 3. Detector window effects on minimum number of photons per pixel

In Sec. 3.3 of the main manuscript document, we performed calculations for determining how many incident photons per pixel $\bar{N}_{\text{inc}}$ are required to detect $\bar{N}_{\text{fluor}}$ x-ray fluorescence photons at a limit of detection (LOD) of $\rho'_{\text{min}} = 0.05$ μg/cm$^2$. We also calculated the associated minimum skin dose $D_{\text{skin}}$ that is then delivered to the incident beam-facing matrix surface. Those calculations involved the forward model outlined Sec. 2 and accounted for the excitation dependence of photoionization partial cross sections (PCSes), Coster-Kronig (CK) transitions, and cascade effects. All of those computations were over ranges of both trace element atomic number $Z$ and single incident photon energy $E_{\text{inc}}$. While we displayed the results for a windowless detector and a vacuum environment in Fig. 1, we show here in Fig. S1 the outcome of exact same calculations for $\bar{N}_{\text{inc}}$, but when accounting for a 25 μm Be window and a $t_g = 5$ cm air gap.

### References


1. R. Jenkins, R. Manne, R. Robin, and C. Senemaud, "Nomenclature, symbols, units and their usage in spectrochemical analysis – VIII. Nomenclature system for x-ray spectroscopy (recommendations 1991)," Pure Appl. Chem. **63**, 735–746 (1991).
2. M. Siegbahn, *The Spectroscopy of X-rays* (Oxford University Press, London, UK, 1925).
3. J. Kirz, D. Sayre, and J. Dilger, "Comparative analysis of x-ray emission microscopies for biological specimens," Ann. New York Acad. Sci. **306**, 291–305 (1978).
4. C. Jacobsen, *X-ray Microscopy* (Cambridge University Press, Cambridge, UK, 2020).
5. B. L. Henke, E. M. Gullikson, and J. C. Davis, "X-ray interactions: Photoabsorption, scattering, transmission, and reflection at $E$=50–30,000 eV, $Z$=1–92," At. Data Nucl. Data Tables **54**, 181–342 (1993).
6. R. D. Deslattes, "Estimates of x-ray attenuation coefficients for the elements and their compounds," Acta Crystallogr. A **25**, 89–93 (1969).
7. E. C. McCullough, "Photon attenuation in computed tomography," Med. Phys. **2**, 307–320 (1975).
8. T. Schoonjans, A. Brunetti, B. Golosio, *et al.*, "The xraylib library for X-ray–matter interactions. Recent developments," Spectrochimica Acta B **66**, 776–784 (2011).

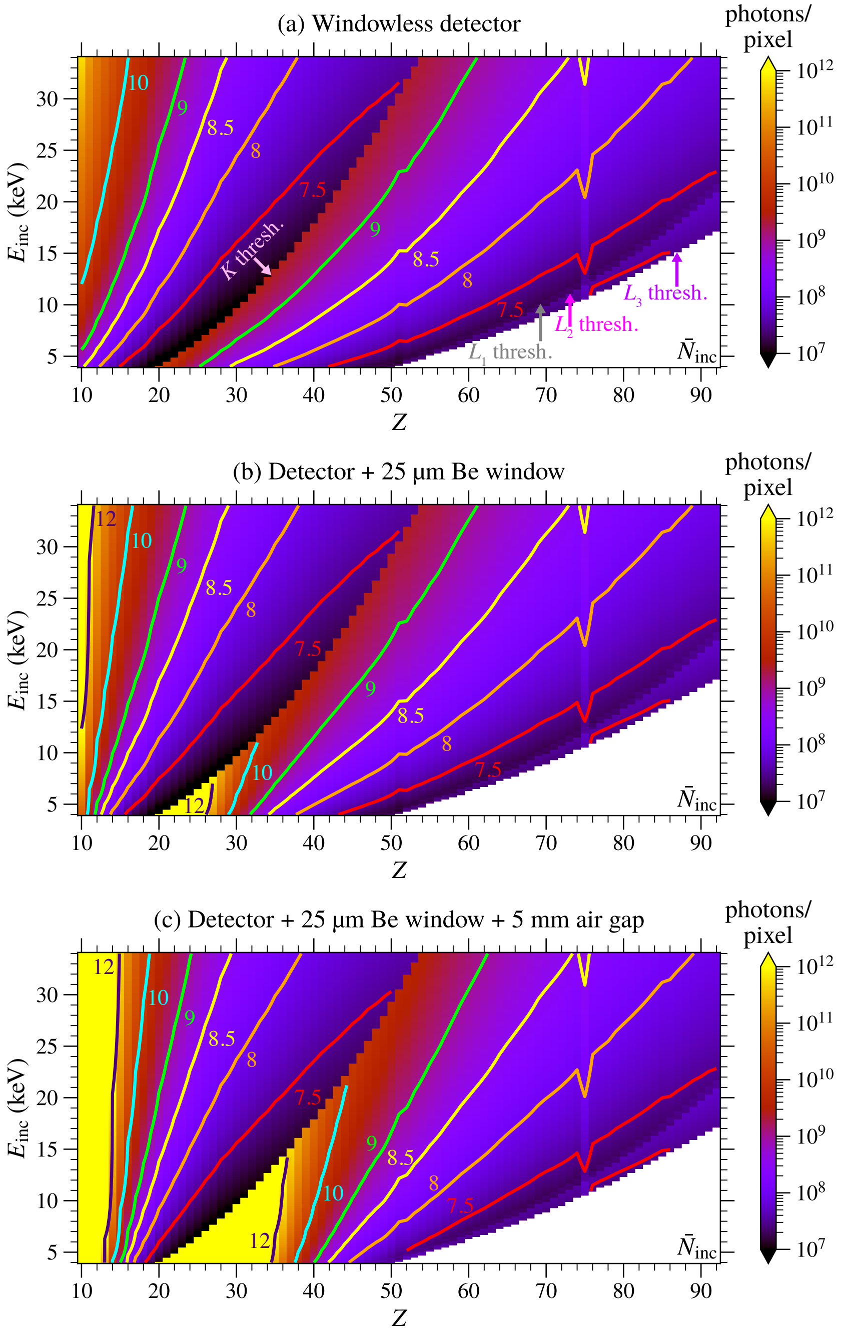


Note: Contour values correspond to base-10 exponents.

Fig. S1. Effects of EDS detector windows and air on the minimum expected number of incident photons $\bar{N}_{inc}$ per pixel required to detect $\bar{N}_{fluor} = 5$ x-ray fluorescence photons per pixel. At top (a) is the combined false color map and contour plot of Fig. 1(a) when neglecting fluorescence signal absorption both in air and in a window mounted onto the front of an energy-dispersive (EDS) detector. At center (b) is the same type of false color map-contour plot combination when accounting for only a 25 µm Be window. The bottom map-contour plot combination (c) shows the effects of both the Be window and a $t_g = 5$ mm air gap between the sample and the detector window. The addition of the window lead to higher $\bar{N}_{inc}$ for lower-$Z$ elements. The inclusion of the air gap further increased $\bar{N}_{inc}$ for an even wider range of low-$Z$ elements. Tabulated data from *xraylib* [8].
